# Identification of DOK2 and PTPN11 as novel interactors of T cell specific adapter protein TSAd

**DOI:** 10.1101/2025.02.11.636929

**Authors:** Hanna Chan, Pawel Borowicz, Brian C Gilmour, Maria Stensland, Ivan Garcia-Loza, Santosh Phuyal, Romina Marone, Lukas T. Jeker, Tuula A Nyman, Anne Spurkland

## Abstract

T cells play a crucial role in the adaptive immune system and depend on tightly regulated intracellular signalling pathways to respond in an appropriate manner. Adapter proteins have flexible and dynamic features, which allow them to regulate T cell signal transduction pathways. As adapter proteins are enzymatically inert and may play multiple roles in parallel, it has been a challenge to fully characterise their functions individually. One such protein in T cells, is T cell specific adapter protein (TSAd), which is upregulated following T cell receptor (TCR) stimulation and is believed to mediate Src family tyrosine kinase signalling. However, the functional role remains elusive, possibly due to limited insight into interactors that potentially bind TSAd. The only structurally well-defined feature within TSAd, is the Src homology 2 (SH2) domain. This conserved domain displays prototypic binding of phosphorylated tyrosines, which suggests that the adapter molecule is implicated in phosphotyrosine signalling pathways. Here, we used an unbiased approach to identify ligands of the TSAd SH2 domain, by using affinity-purification mass spectrometry (AP-MS). Several novel ligands, many of which are known to be implicated in negative regulation of T cell intracellular signalling, were identified. More specifically, we showed that TSAd binds DOK2 and PTPN11 and determined the tyrosines responsible for the TSAd SH2 domain-dependent interaction. Ablation of TSAd and DOK2 by CRISPR/Cas9 in Jurkat T cells resulted in altered tyrosine phosphorylation. Taken together, these findings provide new insight into the possible function of TSAd as a negative signalling node in T cells.

## Introduction

T cells receive stimulatory signals via their T cell receptor (TCR) and surface co-receptors, playing an essential role in adaptive immunity. Incoming cues are transduced along a complex network of signalling pathways. This is largely mediated by phosphorylation cascades, which converge in the cell nucleus resulting in gene expression alterations and consequently T cell activation. For T cells to respond in an appropriate manner to external stimuli, it is imperative that signalling pathways are tightly regulated. A group of molecules, namely adapter proteins, are largely responsible for modulating these signal transduction pathways in a spatiotemporal manner. While adapter proteins are enzymatically inert, they function via their protein binding domains and motifs, harbour structural flexibility and are dynamic. A common binding domain that exists in many adapter proteins of immune cells, is the Src homology 2 (SH2) domain (1).

The SH2 domain is an evolutionary conserved protein structure (2). It consists of approximately 100 amino acids and prototypically binds phosphorylated tyrosines (pTyr) (3). In humans, 121 SH2 domains exist, expressed across 111 proteins (4,5). As such, SH2 domains are essential mediators of pTyr signalling and through protein-protein interactions allow regulation of intracellular signalling.

One of these SH2-domain containing adapter proteins, is T cell specific adapter protein (TSAd), which is upregulated in activated T cells (6) and is believed to act downstream of the TCR signalling cascade. TSAd, encoded by the *SH2D2A* gene, contains an N-terminal SH2 domain followed by several protein interaction motifs, including a proline rich region and multiple conserved tyrosine phosphorylation sites at its C-terminal end (6). As such, it serves as an adapter protein, bridging proteins via its interaction motifs (1).

It has been reported that TSAd SH2 transgenic mice, which overexpressed the isolated TSAd SH2 domain, displayed defects in T cell activation and Th1 mediated immune responses (7).More recently, it has been shown that TSAd-deficient mice display accelerated allograft rejection, suggesting that TSAd mediates regulatory T cell activity (8). However, the underlying molecular mechanisms in which TSAd is implicated in remains unclear, possibly due to the limited overview of TSAd ligands. Given the importance of SH2 domains in pTyr signalling, it is of particular interest to identify ligands of the TSAd SH2 domain to gain a better understanding of its functional role in T cell signalling. The prototypic binding motif for the TSAd SH2 domain has been identified as [H/E/P][pY][D/E/S][N] (9) or [pY][E][N/T][D/ϕ] (10). So far, a few potential interactors in T cells have been suggested, including valosin-containing protein (VCP) (11), and laminin binding protein (LBP) (12). Most recently, using an *in silico* approach, our lab has additionally reported that CD6 and linker of activated T cells (LAT) are potential ligands of the TSAd SH2 domain (13). However, the TSAd SH2 domain interactome has not yet been fully explored and characterised in an unbiased manner.

Here, we identified novel ligands of the TSAd SH2 domain, by using an unbiased approach. Using green fluorescent protein (GFP) tagged TSAd protein, we performed pulldown experiments in Jurkat T cells. Affinity purification mass spectrometry (AP-MS) analysis, revealed potential interaction partners for TSAd SH2 domain. Further, using CRISPR/Cas9 gene editing, we ablated TSAd and one of the newly identified ligands and investigated whether TCR signalling was affected.

## Results

### Novel ligands of the TSAd-SH2 domain identified by affinity-purification mass spectrometry (AP-MS)

Although several ligands of the TSAd-SH2 domain have been identified and suggested in the literature, most have been identified in two-hybrid systems (12,14–16), peptide based arrays (17,18) or using *in silico* methods (13). However, these methods do not consider competitive binding with other proteins. To identify physiologically relevant interactors, we here aimed to identify ligands using an unbiased *in vitro* approach. Initially, using exogenous TSAd-SH2 domain produced in *E. coli*, we performed pulldown using cell lysates from the Jurkat cell line. However, the mass spectrometry (MS) analyses revealed low number of hits (data not shown), possibly due to the inherent challenges during expression of recombinant TSAd-SH2 domain (19). To overcome this issue, we designed two truncated versions of human TSAd with C-terminal tagged GFP for pulldown using anti-GFP nanobodies. The truncated versions include the N-terminal end (TSAd N-term) (1-205a.a. including the SH2 domain (95-187a.a.)) or the C-terminal end (TSAd C-term) (208-389a.a.) (Fig. 1A). As TSAd N-term mainly consists of the SH2 domain, it acted as a surrogate for the recombinant TSAd SH2 domain. The plasmids were transfected into a mutated Jurkat cell line, which displays elevated amounts of pTyr-containing proteins (Jurkat TAg (JTAg) LCK Y192E/Y505F) (20), and thus potentially contain a larger amount of pTyr-containing proteins to which the TSAd SH2 domain could bind. As different cellular stimulation results in different phosphorylation cascades, we treated cells with pervanadate, a tyrosine phosphatase inhibitor (21), to enrich for pTyr-containing proteins (Fig. 1B). Analysis of co-precipitated proteins by immunoblotting, showed that with 2.5 min TCR stimulation, only a limited amount of pTyr-containing proteins were co-precipitated, while pervanadate treatment resulted in increased co-precipitation of pTyr-containing proteins (Fig. S1A).

**Figure 1.**
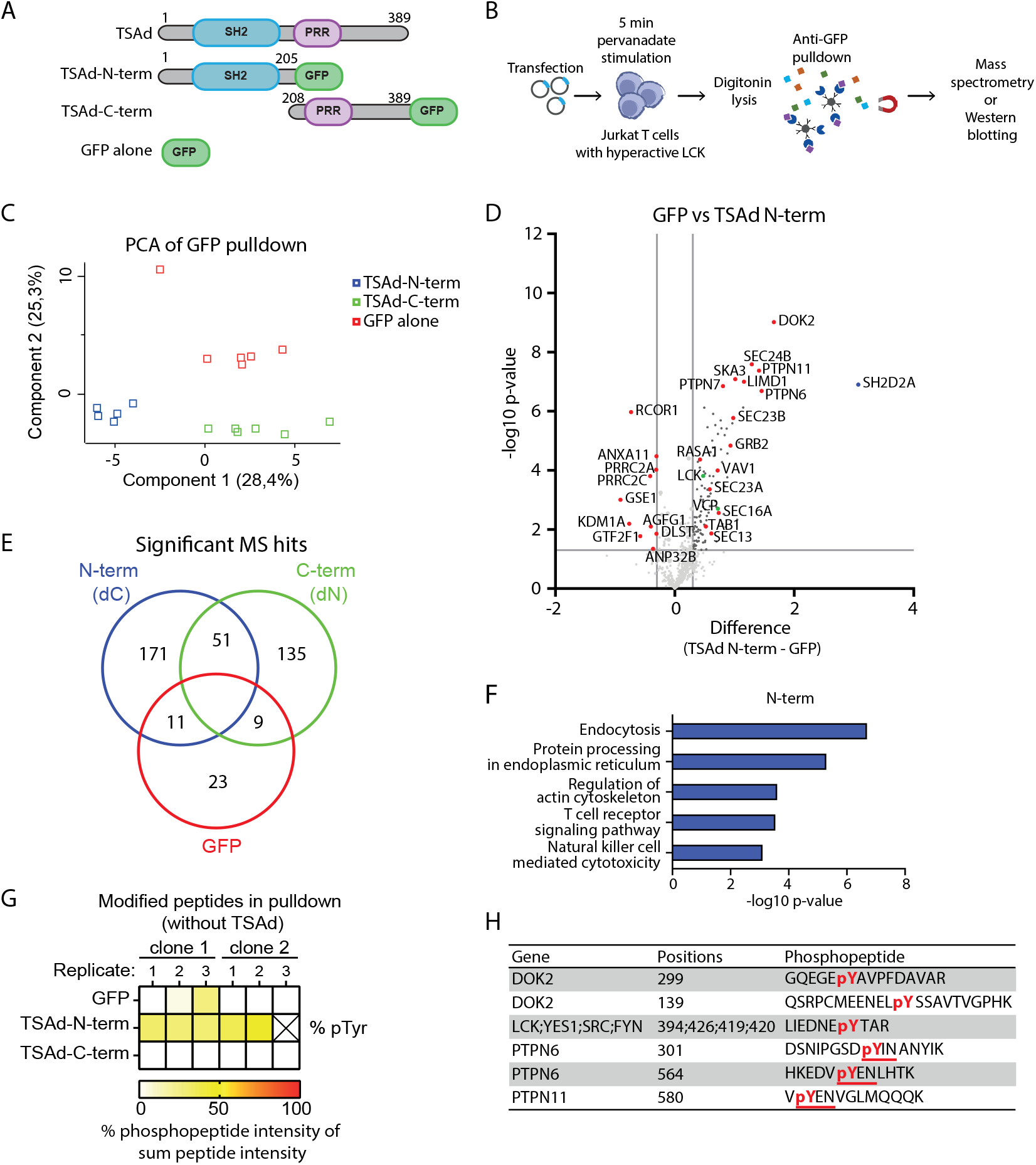
Mass spectrometry analysis identifies ligands of the TSAd-SH2 domain. JTAg Y192E/Y505F cells were transfected with plasmids encoding GFP-tagged TSAd N-term or C-term or GFP alone. 16 hrs after transfection, cells were activated with pervanadate for 5 min, lysed with digitonin lysis buffer and incubated with streptavidin-beads coated with biotin-GFP-nanobodies. Total amount of GFP pulldown sample was split into two; one half was subjected to immunoblotting (Fig. S1A) and the other to MS analysis. A. Schematic of GFP-tagged TSAd constructs. B. Schematic of workflow for GFP-pulldown. C. PCA plot of samples (*n* = 5-6) following analysis of MS results (after removal of values with less than 5 valid hits). Each point represents a replicate of TSAd-N-term (blue), TSAd-C-term (green) or GFP alone (red). D. Volcano plot of the identified hits in GFP vs TSAd-N-term. Proteins enriched in GFP (left) and TSAd-N-term (right). Selected novel interactors marked in red and previously identified ligands marked in green. TSAd (SH2D2A) marked in blue. Horizontal cut-off line denotes p-value = 0.05. Vertical cut-off lines denote fold change = 2. E. Venn diagram of significantly identified proteins in the given samples. Significant hits were identified by first filtering out proteins with at least 5 valid hits among the groups, followed by a two-tailed Student’s T-test . Hits with a p-value<0.05 were determined as significant. F. Bar graph showing enriched KEGG pathways, performed using STRING analysis, with significant hits in TSAd N-term pulldown as identified in (E). G. Heatmap of pTyr-containing phosphopeptides in GFP pulldown analysed by AP-MS. % of phosphopeptide intensity is calculated as the ratio between phosphopeptide intensity versus the total peptide intensity. The “X” in the figure indicates the sample which was removed. H. Table of identified pTyr-containing peptide sequences with a probability score ≥ 0.75 in the TSAd-N-term pulldown. pTyr modification (pY) highlighted in red. pYXN motif underlined in red.

Following GFP pulldown, samples were subjected to MS analyses to identify proteins associated with TSAd (Fig. 1B). Initial principal component analysis (PCA) identified an outlier (Fig. S1B), which we removed from further analysis (described in *Materials and Methods*), and revealed that TSAd N-term and TSAd C-term bind to distinct proteins (Fig. 1C). Focusing on statistically significant novel interactors of TSAd N-term, we identified phosphatases including PTPN6, PTPN7 and PTPN11 and adapter proteins GRB2, TAB1 and DOK2. In addition, a group of proteins implicated in the coat protein complex II (COPII) (22), including SEC24A/B, SEC23B and SEC13 were identified. Of note, previously known ligands of the TSAd SH2 domain were found, including VCP (11) and LCK (23) (Fig. 1D), supporting the design of our approach.

Taking into account all the statistically significant interactors, we identified 171 and 135 hits interacting with TSAd N-term or TSAd C-term respectively (Fig. 1E). Some proteins were found to interact with both ends of TSAd, suggesting presence of multiple docking sites on TSAd. Focusing on the TSAd N-term hits, enriched KEGG pathways include “protein processing in the endoplasmic reticulum” and “TCR signalling pathways” (Fig. 1F), in contrast to pathways identified for TSAd C-term (Fig. S1C-E).

A search for phospho-modifications in the MS dataset revealed that there was no preference for phospho-threonine and –serine modifications, as there was no bias between the TSAd N-term and TSAd C-term GFP-pulldown (Fig. S1F). However, there was enrichment of pTyr-containing peptides among those pulled down by TSAd N-term (Fig. 1G), which was supported by immunoblotting analysis (Fig. S1G), reflecting TSAd SH2 domain pTyr binding activity. In accordance with a preferred binding motif, pYXN, for the TSAd SH2 domain (9), among the pTyr-peptides identified in our MS analysis, several of the hits contained this motif (Fig. 1H). This strongly supports the notion that several of the identified proteins are directly interacting with the TSAd SH2 domain.

### TSAd and its interactors are co-expressed in T cell subsets

One limitation to our approach was the choice of the JTAg cell line for pulldown experiments. This cell line may contain abnormal copy numbers of genes and displays irregular protein expression profiles compared to primary CD4 T cells (24). We thus sought to investigate to what extent TSAd and its identified ligands are co-expressed in primary immune cells. In an effort to explore which proteins are correlated with TSAd expression, we re-analysed (Gilmour et al., in preparation) a publicly available mRNAseq dataset consisting of sequenced PBMCs from healthy controls (27). As TSAd is primarily expressed in T cells (6,25) and natural killer (NK) cells (26), we focused on these subsets in the data (Fig. 2A). Relative to the whole cell population, TSAd is most prominently expressed in NK cells, memory CD8 T cells and MAITs (Fig. 2B).

**Figure 2.**
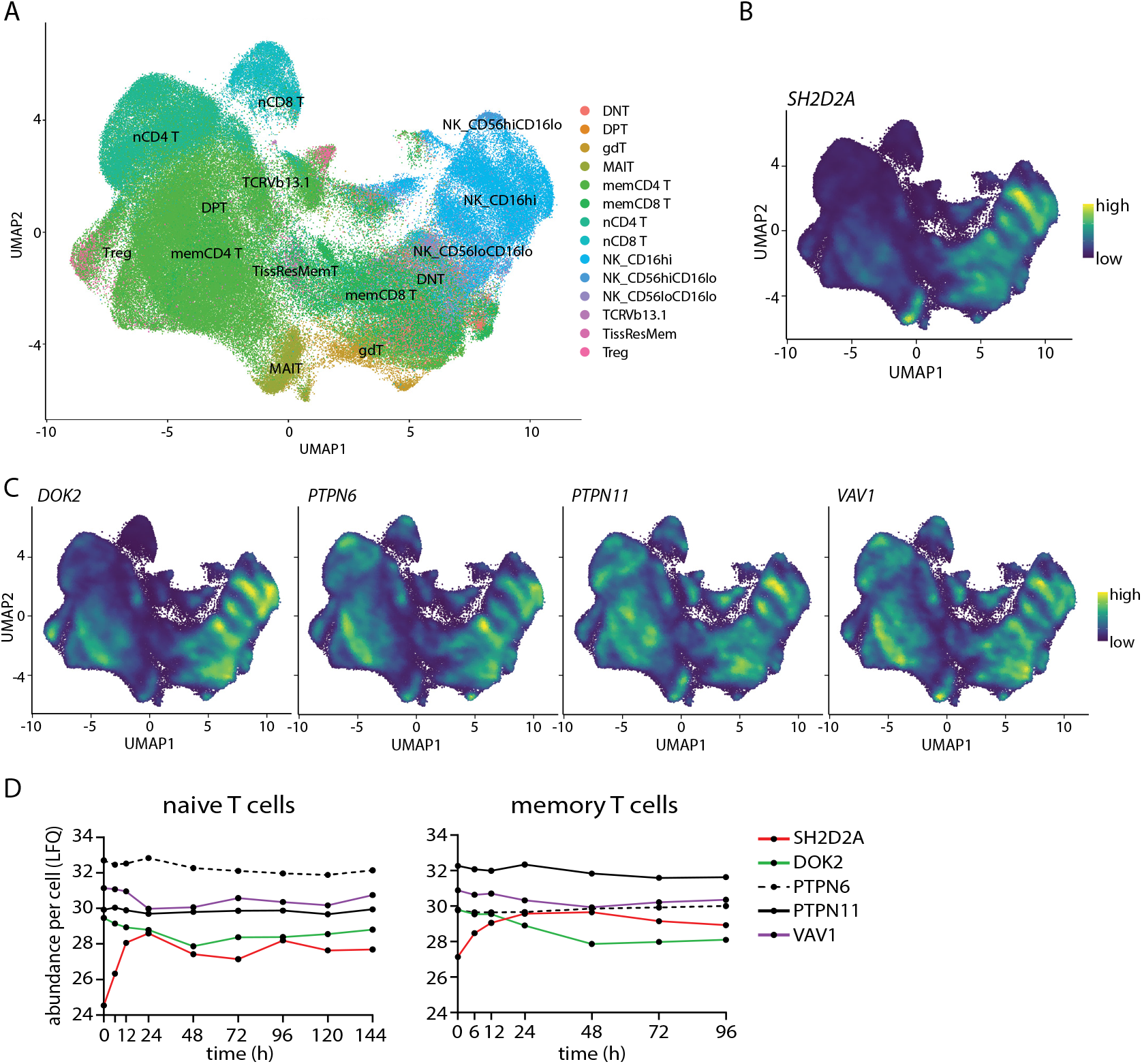
Co-expression of TSAd and its interactors in T and NK cells. A. Transcript-based UMAP visualisation of single cells for T and NK cell subsets, labelled in the corresponding course-level clustering. Data extracted from (27). B-C. Transcript-based UMAP visualisation of T and NK cell types as defined in (A). Clusters are coloured based on expression density from high to low of *SH2D2A* (B), *DOK2, PTPN6, PTPN11* and *VAV1* (C). D. Protein abundance kinetics per cell (LFQ) of SH2D2A, DOK2, PTPN6, PTPN11 and VAV1 from naïve (left) and memory (right) human T cells following anti-TCR stimulation at indicated time points. Data extracted from (33). DNT: double negative T cell; DPT: double positive T cell; gdT: gamma/delta T cell; MAIT: Mucosal-associated invariant T cell; memCD4 T: memory CD4+ T cell; memCD8 T: memory CD8+ T cell; nCD4 T: naïve CD4+ T cell; nCD8 T: naïve CD8+ T cell; NK_CD16hi: natural killer CD16hi; NK_ CD56hiCD16lo: natural killer CD56hiCD16lo; NK_CD56loCD16lo: natural killer CD56loCD16lo; TCRVb13.1: T cell receptor v/beta 13.1 T cell; TissResMemT: tissue resident memory T cell; Treg: regulatory T cell.

Next, we narrowed our focus on the hits identified in the MS analysis, which are known to play regulatory roles in T cell intracellular signalling cascades, including adapter protein DOK2 (28,29), tyrosine phosphatases PTPN6 and PTPN11 (30) and guanine nucleotide exchange factor VAV1 (31). As shown in the UMAP, the adapter protein DOK2 follows a similar expression pattern as TSAd, whereas PTPN6, PTPN11 and VAV1 is ubiquitously expressed across all T and NK cell types (Fig. 2C).

Since mRNA expression of TSAd is not directly correlated with protein expression (32), we next examined how these genes, are expressed on a protein level in T cells. Using publicly available protein data from activated human T cells (33), we observed that TSAd protein is induced by TCR activation in both naïve and memory CD4 T cells (Fig. 2D). In contrast, the potential TSAd interactors maintain a relatively stable protein abundance in naïve cells and are not further induced upon activation. This trend is similar in memory T cells, with the exception of DOK2, which starts to decline 12hrs post-activation. This suggests that the functional roles for DOK2 and TSAd may be different in naïve versus memory T cells. Overall, we show that TSAd and the identified ligands are co-expressed in primary T cells, supporting the notion that TSAd and the identified interactors are likely to be physiologically relevant.

### TSAd interacts with DOK2 and PTPN11 via their C-terminal pTyrs

A conserved arginine in the pTyr binding pocket of SH2 domains largely defines its specificity (34). Mutation of this conserved arginine, disrupts the ability of SH2 domains to bind pTyr motifs. To investigate whether TSAd interactions with DOK2, PTPN6, PTPN11 and VAV1 are dependent on conventional pTyr-SH2 domain binding, we mutated the conserved arginine in our TSAd constructs (TSAd-R120K), repeated the pulldowns and analysed the results by immunoblotting. To avoid interference from endogenous TSAd, the GFP-tagged constructs were expressed in JTAg cells which express wild-type (WT) LCK but not TSAd (i.e. TSAd knockout (KO)).

The results confirmed that DOK2 and PTPN11 interacted with full-length TSAd and TSAd N-term in a pervanadate-dependent manner. Furthermore, we found that the interactions are dependent on the TSAd SH2 domain, as neither TSAd lacking the SH2 domain (TSAd C-term) nor TSAd harbouring the non-functional SH2 domain (R120K mutants) were able to co-precipitate DOK2 or PTPN11 (Fig. 3A-B). In contrast, interactions of TSAd with PTPN6 and VAV1, which interacted with full-length TSAd and TSAd-N-term, were not fully disrupted by lack of a functional SH2 domain (Fig. 3A-B) suggesting multiple binding sites or indirect binding.

**Figure 3.**
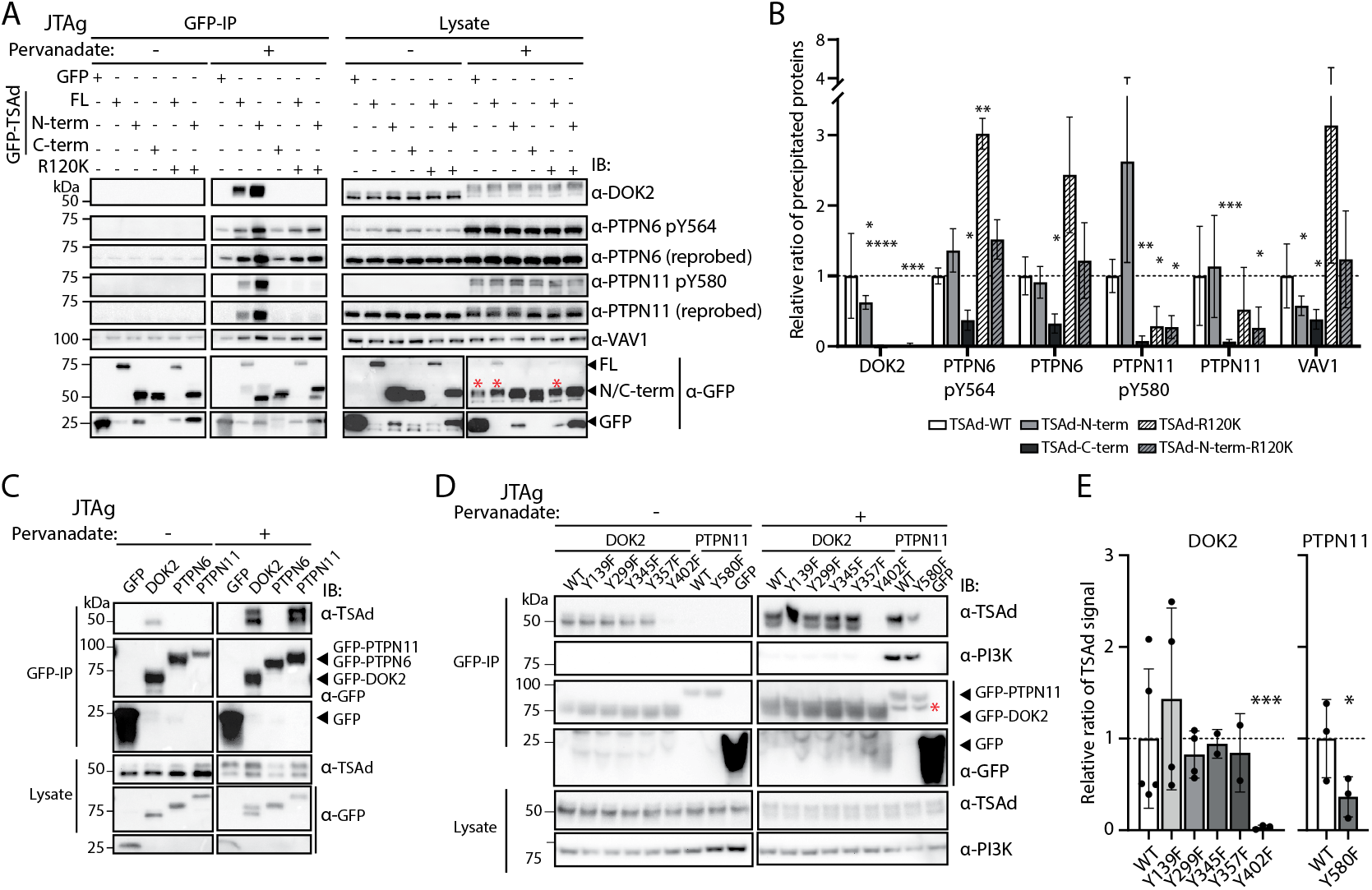
TSAd interacts with DOK2 and PTPN11 in an SH2 domain dependent manner. A. JTAg TSAd KO cells were transfected with GFP-tagged TSAd plasmids. After 36 hrs, cells were left unstimulated or activated for 5 min with pervanadate and lysed with digitonin lysis buffer. Lysates were subjected to GFP pulldown and analysed by immunoblotting. In the lysate blot probed with anti-GFP, bands labelled with a red * above, denotes endogenous DOK2 from the previous probing. One representative immunoblot is shown (*n* = 2-3). FL=full-length construct. IB=immunoblot. B. Bar graph shows quantification of immunoblotting from A, where signals are normalised to the construct’s GFP signal. The average of the WT TSAd was set to 1 and statistical significance was determined with a one-sample t-test (versus value of 1) (*n* = 2-3) C. JTAg cells were transfected with plasmids encoding GFP tagged DOK2, PTPN6 or PTPN11. 5 hrs after transfection, cells were activated with PMA/IO for 16-18 hrs. Cells were rested overnight, then left unstimulated or activated for 5 min with pervanadate and lysed with digitonin lysis buffer. Lysates were subjected to GFP pulldown and analysed by immunoblotting. D-E. Same as in C, but here JTAg cells were transfected with plasmids encoding GFP tagged DOK2 or PTPN11 mutants. A representative immunoblot is shown (D) and the bar graph shows the quantification where signals are normalised to the construct’s GFP signal. Average of the WT construct was set to 1 and statistical significance was determined with a one-sample t-test (versus value of 1) then to WT TSAd sample (E). (*n* = 3-4). In the GFP-IP blot probed with anti-GFP, the second band labelled with a red * in the pervanadate stimulated PTPN11 sample, is the PI3K p85α-subunit band from the previous probing. All bar graphs show mean with SD.

We then performed the pulldown in reverse, and pulled down GFP-tagged DOK2, PTPN6 and PTPN11 and probed for co-precipitation of endogenously expressed TSAd (Fig. 3C). In unstimulated cells, interaction of PTPN6 nor PTPN11 to TSAd was observed, while TSAd was slightly associated with DOK2. In cells treated with pervanadate, both DOK2 and PTPN11, but not PTPN6 bound to TSAd. Taken together, our findings suggest that TSAd interacts with both DOK2 and PTPN11 in a pTyr-dependent manner. In contrast, while PTPN6 was initially a candidate identified in the MS dataset, the reverse pulldown indicated that it most likely does not directly interact with TSAd.

Having found that TSAd bound to DOK2 and PTPN11 in a pTyr dependent manner, we aimed to identify the tyrosines within DOK2 and PTPN11 that were responsible for the interactions. Mutation of Tyr to Phe within SH2 domain binding motifs, mimics the non-phosphorylated form of Tyr (35), thus destroying the interaction site for an SH2-domain. To this end, we mutated the pTyr sites identified in our MS dataset (Fig. 1H). In addition, as the canonical motif for TSAd was not found among the DOK2 phosphopeptides in our MS dataset, we mutated several additional tyrosines in DOK2 reported by others to be commonly phosphorylated (36). Pulldowns from lysates of Jurkat cells expressing these mutants revealed that phosphorylated DOK2-Tyr402 and PTPN11-Tyr580 are essential for interaction of TSAd with DOK2 and PTPN11 respectively (Fig. 3D-E), strongly suggesting that these tyrosines represent binding motifs for the TSAd SH2 domain. As PTPN11 and the p85 subunit of phosphoinositide 3-kinase (PI3K) are known to interact in a p85-subunit-SH2 domain dependent manner (37), we probed for PI3K to ascertain that the PTPN11 Phe580 mutation did not disrupt this binding. Both WT PTPN11 and PTPN11 Y580F bound to PI3K (Fig. 3D), confirming that the PI3K is binding to PTPN11 independent of Tyr580 phosphorylation, in contrast to what we observe with TSAd.

### TSAd interacts with DOK2 and PTPN11 in live cells

Although pulldown assays aid in identifying protein-protein interactions, they do not take into consideration the spatial component. In the lysates, proteins may be interacting after being released from their respective subcellular locations. To investigate whether and where binding of TSAd to DOK2 or PTPN11 occur within live cells, we used the bimolecular fluorescence complementation (BiFC) assay (38,39). This approach relies on the interaction between two halves of a yellow fluorescent protein (YFP), the N-terminal and C-terminal YFP termed as nYFP and cYFP respectively. When interactions between the co-transfected fusion proteins (Fig. S2) occur, the nYFP and cYFP are brought into the vicinity of each other and form a mature YFP protein, which emits a fluorescent signal (Fig. 4A). DOK2 is a cytosolic protein, which has a pleckstrin homology (PH) domain, allowing it to be recruited to the cell membrane via binding to PI(3,4,5)P3 (40), which is normally upregulated in activated T cells (41). However, in JTAg cells, PI(3,4,5)P3 is constitutively present due to the lack of SHIP1 and PTEN lipid phosphatase expression (41,42). Given these properties, TSAd and DOK2 were found to interact at the cell membrane in JTAg cells (Fig. 4B). Consistent with the pulldown results (Fig. 3A), the interaction was affected by the TSAd R120K mutation, where the dot-like pattern at the membrane disappeared and the signal was more diffused in the cell. PTPN11 is also cytosolic, but unlike DOK2, it does not contain a PH domain nor other membrane localisation features. We found TSAd-bound PTPN11 spread out in the cytosol of JTAg cells and the interaction was disrupted by the TSAd-R120K mutation (Fig. 4B), in line with the pulldown experiments (Fig. 3D-E). Taken together, these observations further verify the hits found in the MS analysis and suggest possible cellular locations of these interactions.

**Figure 4.**
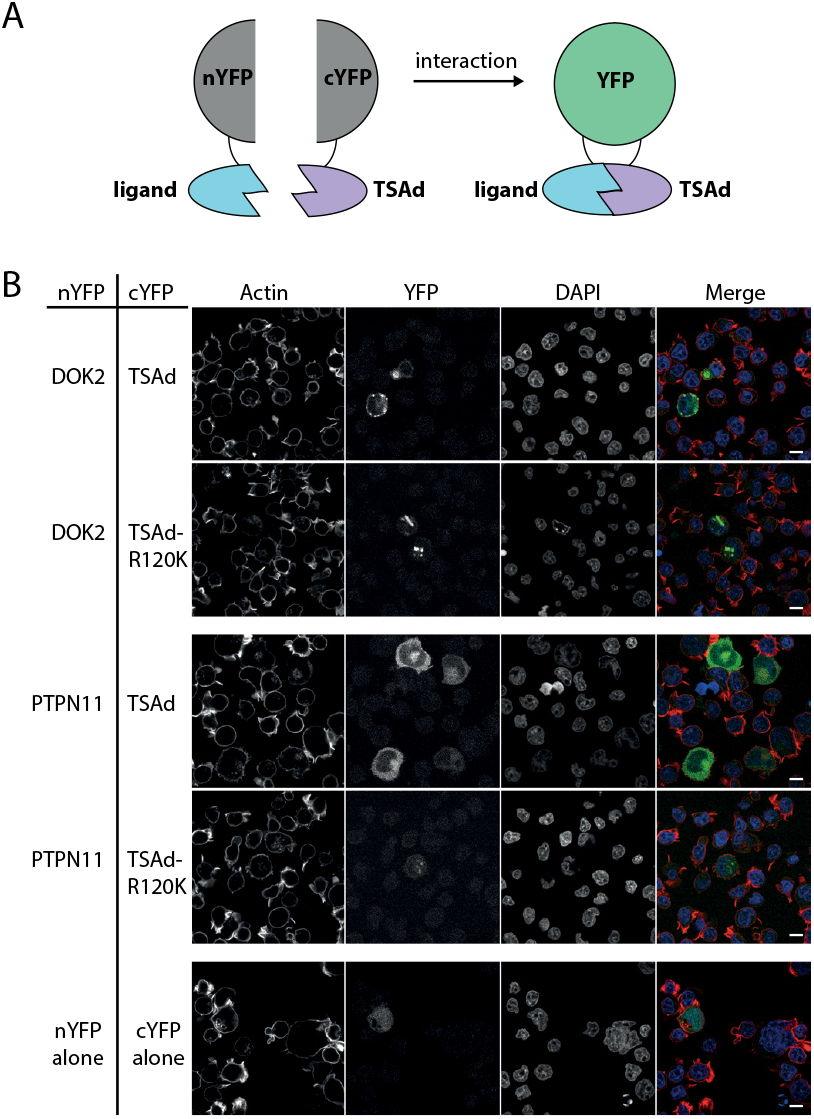
TSAd interacts with DOK2 and PTPN11 in live cells. A. Schematic of the BiFC assay. B. JTAg TSAd KO cells were transfected with plasmids encoding nYFP and cYFP tagged constructs. After 48 hrs, cells were activated with pervanadate for 5 min, washed, and incubated for 4 hrs. Cells were then prepared for imaging and stained for nucleus and actin. Representative images from the BiFC between TSAd and DOK2, and TSAd and PTPN11 are shown. Co-transfected constructs are noted in the table to the left of the image. Scale bar = 10μM. Merged image of actin (red), YFP (green) and nucleus (blue). Lysates from cells transfected with DOK2 and PTPN11 were subjected to immunoblotting in Fig. S2.

### Ablation of TSAd and/or DOK2 results in elevated pTyr events

As DOK2 has been reported to be a negative regulator of TCR signalling (28,29), we next investigated the effects of phosphorylation kinetics downstream of TCR activation in the presence or absence of TSAd and/or DOK2. To this end, using CRISPR/Cas9 gene editing, we generated TSAd KO, DOK2 KO or double KO Jurkat E6.1 cells (Fig. S3A). Compared to JTAg cells, Jurkat E6.1 cells have a phenotype closer to that of non-transformed T cells (20), hence are more appropriate cells to study T cell phenotypes. Following overnight serum starvation of Jurkat E6.1 cells, we monitored pTyr-kinetics downstream of TCR stimulation. Compared to wild-type (WT) cells, there was a slightly elevated total pTyr signal in both TSAd KO and DOK2 KO both at basal levels and following TCR stimulation. However, in the double KO cells the total pTyr signal was reduced compared to WT cells (Fig. 5A-B). Taken together, these observations suggest a potential inhibitory role of TSAd and DOK2 as protein interactors.

**Figure 5.**
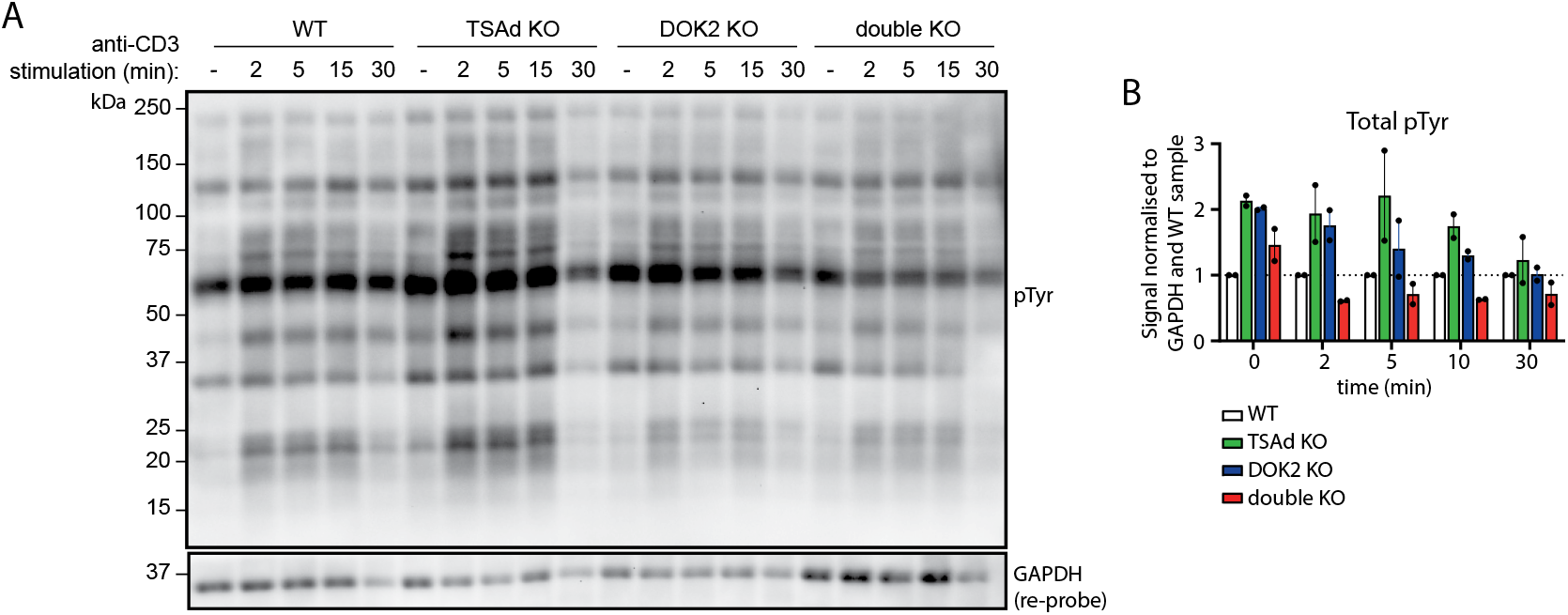
Lack of TSAd or DOK2 results in increased pTyr events, but does not disrupt proximal TCR signalling. TSAd alone, DOK2 alone or both (double KO) were knocked-out in Jurkat E6.1 cells by CRISPR/Cas9 gene editing and phosphorTyr (pTyr) kinetics were monitored. A. Gene edited or WT Jurkat E6.1 cells were serum-starved overnight and left unstimulated (0 min) or stimulated with anti-CD3ε antibodies for 2, 5, 15 or 30 min as indicated. Lysates were resolved on an immunoblot and pTyr (A) was probed for. In 5A, GAPDH was re-probed after pTyr probing. One representative blot from two biological replicates is shown. B. Quantification of pTyr kinetic experiments from A (*n* = 2, from two independent experiments). Bar graph shows median with range.

## Discussion

Since the discovery of TSAd, several groups have identified and studied the functional role of TSAd in T cells. As TSAd is upregulated following T cell activation and has high protein conservation across species (1,6), it has been considered a possible mediator of signalling in experienced T cells. However, TSAd appears dispensable, making it a challenge to characterise its defined function in T cells. Given its structural properties, TSAd most likely exerts its functions by bridging proteins. As the only structurally defined feature of TSAd is its SH2 domain, mapping out the interactome of the TSAd SH2 domain is crucial to elucidate the protein’s function. Although efforts have been made to identify and characterise ligands of the TSAd SH2 domain, the approaches taken so far have been biased, in form of peptide arrays (9,11,12,43) or using *in silico* approaches (13). To our knowledge, this is the first report of using an unbiased approach *in vitro* to identify ligands of the TSAd SH2 domain. Using GFP-tagged constructs of truncated TSAd we performed AP-MS and identified cytosolic ligands of N-terminal TSAd (containing the SH2 domain) in the Jurkat TAg cell line. Our MS analysis identified several novel TSAd interactors, with unique functions, suggesting multiple functional roles for TSAd.

Using biochemical methods and reverse genetics, we showed that DOK2 and PTPN11 bind to TSAd in a SH2 domain-dependent manner, where pTyr402 in DOK2 and pTyr580 in PTPN11 were required for the interaction. In contrast, PTPN6 and VAV1 interaction with the TSAd SH2 domain could not be verified by pulldown and immunoblotting analysis, possibly due to indirect binding to TSAd. This highlights the importance of verifying protein-protein interactions identified in MS by other means.

Similar to TSAd, DOK2 is a cytosolic adapter protein and belongs to the DOK protein family (44). DOK2, which is preferentially expressed in hematopoietic cells, contains an N-terminal PH domain followed by a phosphotyrosine binding (PTB) domain and several tyrosines that can be phosphorylated and recognised by SH2 domains (44,45). The function of DOK2 has largely been studied in T cells. It has been suggested as a negative regulator in the context of T cell signalling, by recruiting inhibitory effectors, including SHIP1 and CSK, to the membrane bound Src kinases (28,46). In line with this, we observed elevated total pTyr signals in TSAd KO and DOK2 KO cells, which were ameliorated in double KOs. This could be a result of the inhibitory effectors that DOK2 has been reported to interact with, including the negative regulator CSK (47–49). Although we observe altered pTyr kinetics in the absence of TSAd or DOK2 in Jurkat E6.1 cells, it lacks SHIP1 (50), a ligand of DOK2, that may be important in regulating TCR signalling (51). Consequently, what the functional relevance of the TSAd-DOK2 interaction is, remains to be addressed, preferably in primary T cells.

PTPN11 was another interactor of the TSAd SH2 domain verified in this study. PTPN11, also known as SHP-2, is a ubiquitously expressed phosphatase, which contains two SH2 domains in tandem followed by the protein-tyrosine phosphatase (PTP) domain. Its activity is controlled by binding of the N-SH2 domain to the phosphatase core in the inactive state, preventing ligand interaction with the phosphatase. The inhibition is relieved when the SH2 domain binds to pTyr-substrates. In T cells, PTPN11 acts downstream of cytokine receptors as well as inhibitory receptors including PD-1 in the context of T cell exhaustion (52). We found that the Tyr580 in PTPN11 is a binding site for the TSAd SH2 domain. Other ligands of pTyr580 include Grb2 (53) and Fyn (54). In the case of Grb2, it has been shown to act as a *trans*-regulator of PTPN11 function by interacting with PTPN11-pTyr580 via its SH2 domain (55). As the Grb2 SH2 domain and TSAd SH2 domain are predicted to recognise similar pTyr motifs (9), it is possible that TSAd regulates PTPN11 function in a similar or competitive manner.

TSAd has mainly been studied in the context of TCR stimulation and although both DOK2 and PTPN11 may act downstream of TCR activation, they have also been found to be implicated downstream of other surface receptors. Both CD2 and epidermal growth factor receptor (EGFR) can activate DOK2 (48,56), and by inference they could be considered as alternative receptor activators of TSAd. As such, further investigation is needed to identify which stimulants of these receptors are able to induce TSAd and DOK2 or PTPN11 interactions . Moreover, this may reveal the potential roles for TSAd downstream of these signalling pathways. As the lysis method in this study mainly isolated the cytosolic fraction of the cells, it remains unclear if TSAd can exert its function at the membrane by directly interacting with membrane-bound proteins or if TSAd needs to be recruited via membrane-recruiting proteins, such as DOK2. In addition, our co-expression analysis showed that TSAd and its ligands have varied patterns of co-expression across T and NK cells. Investigating the molecular interactions in other T cell subsets or NK cells may provide further insight into their functional roles.

In conclusion, the unbiased approach used in this study allowed us to identify novel ligands of the TSAd SH2 domain, expanding the current overview of TSAd’s interactome in T cells. However, further research is necessary to fully comprehend the functional significance of these interactions, particularly in relation to DOK2 and PTPN11. We are currently investigating the phenotypes of primary T cells deficient in TSAd and/or its ligands, utilising physiological methods of T cell stimulation. Taken together, these efforts may provide new insights into the otherwise elusive function of TSAd in T cells.

## Methods and Materials

### Antibodies and reagents

The following primary antibodies were used: anti-DOK2 (clone E-10, Santa Cruz), anti-pTyr564 PTPN6 (clone D11G5, Cell Signaling Technology), anti-PTPN6 (clone C-19, Santa Cruz), anti-pTyr580 PTPN11 (#3703, Cell Signaling Technology), anti-PTPN11 (clone B-1, Santa Cruz), anti-VAV1 (clone C-14, Santa Cruz Biotechnology), anti-TSAd (clone OTI3C7, Origene), anti-PI3 Kinase p85α subunit (clone AB6, Upstate), anti-GFP (clone B-2, Santa Cruz), anti-pTyr (clone 4G10, Upstate Biotechnology), and anti-GADPH (clone 6C5, Chemicon). The following secondary antibodies were used: HRP conjugated goat anti-mouse (H+L) IgG and HRP conjugated mouse anti-rabbit IgG (Jackson ImmunoResearch). The following reagents were used for stimulation of cells: anti-CD3ε (clone OKT-3, American Type Culture Collection), phorbol 12-myristate 13-acetate (PMA), ionomycin (IO), sodium orthovanadate and hydrogen peroxide (pervanadate) (all Sigma-Aldrich ).

### Plasmids

The GFP-tagged human TSAd construct was generated in the peGFP-N1 vector (Clontech). Restriction enzyme cloning was used to generate GFP-tagged TSAd mutants, where TSAd-N-term consists of human TSAd 1-205aa, which contains the SH2 domain (95-187aa) and the TSAd-C-term construct consisting of 208-389aa. GFP-tagged DOK2 and PTPN11 were cloned from a cDNA library into the GFP vector by Gibson Assembly (New England Biolabs and (57)). GFP tagged PTPN6 was cloned from PTPN6-containing plasmid (a kind gift from Prof. Hans-Christian Åsheim) into the GFP vector by Gibson Assembly. Point mutations were introduced into the plasmids by QuickChange mutagenesis using Pfu-Turbo DNA polymerase (Agilent). All constructs were verified by Sanger sequencing (GATC Eurofins). Split YFP tagged constructs were cloned from the nYFP and cYFP vectors (a kind gift from Prof. Hesso Farhan) using Gibson Assembly.

### Cells and transfection

Jurkat TAg cells WT and mutants, and Jurkat E6.1 WT and mutants, were cultured in RPMI-1640 supplemented with 10% fetal calf serum (FCS), 1mM sodium pyruvate, 1mM non-essential amino acids, 1mM HEPES buffer, 100units/ml penicillin and 100μg/ml streptomycin (all from Gibco-BRL, ThermoFisher Scientific) and 50μM β-mercaptoethanol (Sigma-Aldrich). Jurkat TAg cells were subcloned as described in (1) and all experiments were repeated in at least two independent clonal cell lines with the respective genetic mutations. Jurkat TAg Lck Y192E/Y505F mutants and Jurkat TAg TSAd knockouts were previously described (1). JTAg cell line transfections were performed using a BTX electroporator (Genetronix) at 240 V and 25 ms. Transient transfectants were cultured for 16–24h or as otherwise indicated.

### Stimulation of cells

For overnight (16-18hrs) PMA/IO stimulation, cells were stimulated with 50 ng/ml PMA and 500 ng/ml IO at a concentration of up to 1×106 cells/ml in complete RPMI. For pervanadate treatment, cells were suspended in PBS at a concentration of up to 100×106 cells/ml and pre-warmed for 5 min at 37°C in a water bath. Then, a mix of 0.01 mM sodium orthovanadate (Sigma-Aldrich) and 0.01% hydrogen peroxide (Sigma-Aldrich) was added, and cells were activated for 5 min at 37°C. Treatment was stopped by diluting the samples 10X with cold PBS. For TCR stimulation, cells were suspended in PBS at a concentration of up to 100×106 cells/ml and pre-warmed for 5 min at 37°C in a water bath. Cells were stimulated with 5 μg/ml anti-CD3ε antibody (OKT3) for the indicated time. For TCR activated pTyr-kinetics experiments in Fig. 5, Jurkat E6.1 cells were serum starved for 18 hrs prior to TCR stimulation. Stimulation was stopped by moving a fraction of the sample from the stimulation tube to another tube with cold PBS, diluting the sample 10X, at the indicated time points.

### GFP immunoprecipitation

Samples that were subjected to MS analysis were prepared as follows. 10×106 Jurkat TAg cells harbouring the Lck Y192E/Y505F mutations were transfected with 8μg TSAd-GFP-N-term, 20μg TSAd-GFP-C-term or 5μg GFP alone, in the presence of a spike of 2μg HA-TSAd-Nterm in all samples as this has previously been observed by our group to increase the protein expression of exogenous TSAd constructs (data not shown). Two Jurkat Lck Y192E/Y505F clones were included, and three technical replicates were performed for each clone and for each transfected construct.

After 16 hrs incubation, cells were subjected to pervanadate stimulation as described above. Cells were subsequently subjected to digitonin lysis containing 150mM NaCl, 50mM HEPES (pH7.4), 1X protease inhibitor cocktail (Sigma), 1mM sodium orthovanadate (Sigma Aldrich), 25mM NaF (Fluka), 150μg/ml digitonin (Calbiochem) for 10 min at 4°C on a rotating wheel, before being spun down at 2000xg, 10 min at 4°C. Lysates were incubated with streptavidin coated beads (ThermoFisher), covered with biotinylated anti-GFP VhH nanobodies (Chromotek) for 1 hr with constant rotation. Beads were then washed and subjected to mass spectrometry analysis or analysed by immunoblotting. All replicates were processed and analysed by mass spectrometry in the same experiment.

### Immunoblotting

Lysates or pulldown beads were denatured in SDS loading buffer containing 0.35M Tris-HCl, 10% SDS, 6% β-mercaptoethanol (Sigma-Aldrich), 30% glycerol (VWR international S.A.S), 0.175mM bromophenol blue (Fluka Ag), pH 6.8), and boiled at 95°C for 10min. Denatured proteins were resolved by SDS-PAGE and transferred to PVDF membranes with the Trans-Blot Turbo Transfer System (all Bio-Rad laboratories). Where lysates were resolved on a SDS-PAGE gel, 2.5% of the input was resolved. The membranes were blocked and incubated overnight at 4°C with primary antibodies diluted in tris-buffered saline (pH 7.4), 0,1% Tween (Sigma-Aldrich), and 3% skimmed milk or 3% bovine serum albumin (Bio-Rad laboratories) (the latter in combination with anti-pTyr antibodies). Membranes were incubated with the appropriate HRP-conjugated secondary antibody for 1 hr at room temperature. SuperSignal West Pico Stable Peroxide Solution (Pierce) was used to visualise bands using the ChemiDoc Imaging System (Bio-Rad laboratories). Images were analysed with ImageJ (version 1.52a).

### Protein on-beads digestion for mass spectrometry

Beads were resuspended in 10µl 0.2% ProteaseMax Surfactant (Promega) in 50mM NH4HCO3 (Sigma Aldrich) and were further diluted in 100µl 50mM NH4HCO3. Cysteines were reduced in 5mM DTT (Sigma Aldrich) at 56°C for 30 min and alkylated in 15mM Iodoacetamide (Sigma Aldrich) for 20 min at room temperature before overnight digestion by trypsin (1µg) at 37°C (Promega). The resulting peptides were desalted by home-made STAGE tip C18 columns prepared by stacking three layers of C18 Empore Extraction Disk (Varian) into 200μL pipette tips.

### Mass spectrometry

Desalted peptide samples were analyzed by Easy nLC1000 nano-LC system connected to a quadrupole – Orbitrap (QExactive) mass spectrometer (ThermoElectron) equipped with a nanoelectrospray ion source (EasySpray/ Thermo). For liquid chromatography separation, we used an EasySpray column (C18, 2 µm beads, 100 Å, 75 μm inner diameter) (Thermo) capillary of 25 cm bed length. The flow rate used was 0.3 μL/min, and the solvent gradient was 2% to 7 % for 5 min and then to 30% B for 60 min. The column was finally washed in 90 % B wash for 20 min. Solvent A was aqueous 0.1 % formic acid, whereas solvent B was 100 % acetonitrile in 0.1 % formic acid. Column temperature was kept at 60°C.

The mass spectrometer was operated in the data-dependent mode to automatically switch between MS and MS/MS acquisition. Survey full scan MS spectra (from m/z 300 to 1,750) were acquired in the Orbitrap with resolution R = 60,000 at m/z 200 (after accumulation to a target of 3,000,000 ions in the quadruple). The method used allowed sequential isolation of the most intense multiply-charged ions, up to ten, depending on signal intensity, for fragmentation on the HCD cell using high-energy collision dissociation at a target value of 100,000 charges or maximum acquisition time of 20 ms. MS/ MS scans were collected at 15,000 resolution at the Orbitrap cell. Target ions already selected for MS/MS were dynamically excluded for 30 s. General mass spectrometry conditions were: electrospray voltage set to 2.1 kV, no sheath and auxiliary gas flow, heated capillary temperature of 250oC, normalised HCD collision energy 25%.

### Protein identification and label-free quantitation

MS raw files were submitted to MaxQuant software version 1.6.17.0 (58) for protein identification and quantification. Parameters were set as follows: carbamidomethylation as fixed modification and protein N-acetylation, methionine oxidation and phosphorylation (STY) as variable modifications. First search error window of 20 ppm and main search error of 4.5 ppm. Trypsin without proline restriction enzyme option was used, with two allowed miscleavages. Minimal unique peptides were set to 1, and FDR allowed was 0.01 (1%) for peptide and protein identification. The UniProt human reviewed database was used (access date 2020) with GFP added. Generation of reversed sequences was selected to assign FDR rates. Perseus ver 1.6.1.3 was used for the statistical analysis of MaxQuant results. PCA plot of the associated proteins identified by mass spectrometry, revealed that one of the technical replicates (TSAd-Nterm, clone 2) was an outlier and grouped with the GFP control samples (Fig. S1B). Thus, this replicate was removed from subsequent data analysis.

### Bimolecular fluorescence complementation (BiFC) and image analysis

After transfection of the split YFP tagged constructs, cells were activated for 5 min with pervanadate, washed and incubated for 4 hrs to allow for YFP protein maturation. Subsequently, cells were adhered to microscope glass slides (VWR) by incubating at 37°C, for 45 min in PBS. Cells were fixed with 4% PFA (Sigma-Aldrich) in PBS for 15 min, washed twice with PBS, and permeabilised with 0.1% saponin in PBS (PBS-S) for 10 min at room temperature. Cells were stained with DAPI (Invitrogen, Thermo Fisher Scientific), and phalloidin-AF647 (Invitrogen, Thermo Fisher Scientific) in PBS-S for 15 min and washed twice with PBS-S. SlowFade Gold (Life Technologies) was added to each sample, before the glass slide was covered with a cover slip and sealed with nail polish. Samples were stored at -20°C before acquisition of confocal images on the Zeiss LSM-710 Confocal Microscope. Images were analysed by ImageJ (version 1.53t).

### Analysis of public datasets

Publicly available mRNA single sequencing data (27) from a cohort of COVID-19 patients and age-matched controls, was used to extract information on gene expression profiles with single cell resolution. Analysis with mRNA single sequencing data was performed in R using the library Seurat v4.0. Protein expression data were extracted from publicly available data from http://www.immunomics.ch (33), where freshly isolated and sorted naïve or memory CD4+ T cells from human PBMCs were activated and protein abundance over time was monitored by mass spectrometry.

### CRISPR/Cas9 editing

Cas9 ribonucleoproteins (RNPs) were generated based on a previously described method (59) and either generated immediately prior to experiment or used after storage at -80°C. Synthetic crRNAs and tracrRNA (IDT) were resuspended in IDTE buffer to a final concentration of 200μM and stored in aliquots at −80°C. crRNA sequence targeting SH2D2A is 5’-CCAGAACCTGGGCTACACTG-3’ and DOK2 is 5’-GGAGACGGGGCAGTGAAACA-3’. gRNAs were generated by mixing equimolar amounts of crRNA and tracrRNA to a final duplex concentration of 44μM. The duplex was annealed at 95°C for 5 min, then left to cool down at room temperature. To improve editing efficiency, 100mg/ml poly-L-glutamic acid (PGA) diluted in dH2O (Sigma) was added at a 0.8:1 volume ratio prior to adding 1:1 volume ratio of Cas9 enzyme (38μM stock) (University of Geneve Protein Platform) (60). The Cas9 RNP was incubated at room temperature for 30 min. Prior to electroporation, 0.5×106 Jurkat E6.1 cells were washed with PBS, then mixed with 9μl of Neon buffer R (Thermo Fisher Scientific) and 1μl of Cas9 RNPs. Cells were electroporated with the Neon transfection system (Thermo Fisher Scientific) using the 10μl NeonTip with the settings: 1600V, 10 ms pulse width, 3 pulses. Electroporated cells were immediately seeded into complete RPMI and expanded. Editing efficiency was verified by TIDE analysis (61) and immunoblotting (Fig. S3).

### Statistical analysis

GraphPad Prism (v. 9.4.1) was used to perform statistical analysis. Statistical significance of quantified and normalised Western blots was determined using a one-sample t-test (versus value of 1). All statistically significant comparisons are shown on the graphs accordingly: *p<0.05, **p<0.01, ***p<0.001, ****p<0.0001.

## Author contributions

A.S. and H.C. conceptualised the study. H.C., B.C.G. and A.S. designed experiments and analysed results. H.C. performed pulldowns, immunoblotting, molecular cloning of constructs, BiFC assay and CRISPR/Cas9 knockout experiments in Jurkat E6.1 cell lines. P.B. designed experiments, generated JTAg mutants and assisted with data interpretation. H.C. and I.G.L. created YFP and GFP tagged DOK2, PTPN6 and PTPN11 plasmids. B.C.G. analysed publicly available RNA-seq datasets. S.P. provided BiFC reagents and advice. M.S. and T.A.N. analysed mass spectrometry data. R.M. and L.T.J. provided assistance with design of CRISPR/ Cas9 experiments and reagents. H.C. wrote the manuscript with input from all co-authors. A.S. oversaw the project and reviewed the final manuscript.

## Acknowledgments

Mass spectrometry-based proteomic analyses were performed by the Proteomics Core Facility, Department of Immunology, and University of Oslo/Oslo University Hospital, which is supported by the Core Facilities program of the South-Eastern Norway Regional Health Authority. This core facility is also a member of the National Network of Advanced Proteomics Infrastructure (NAPI), which is funded by the Research Council of Norway INFRASTRUKTUR-program (project number: 295910).

