## Supplementary figures for "Identification of DOK2 and PTPN11 as novel interactors of T cell specific adapter protein TSAd"

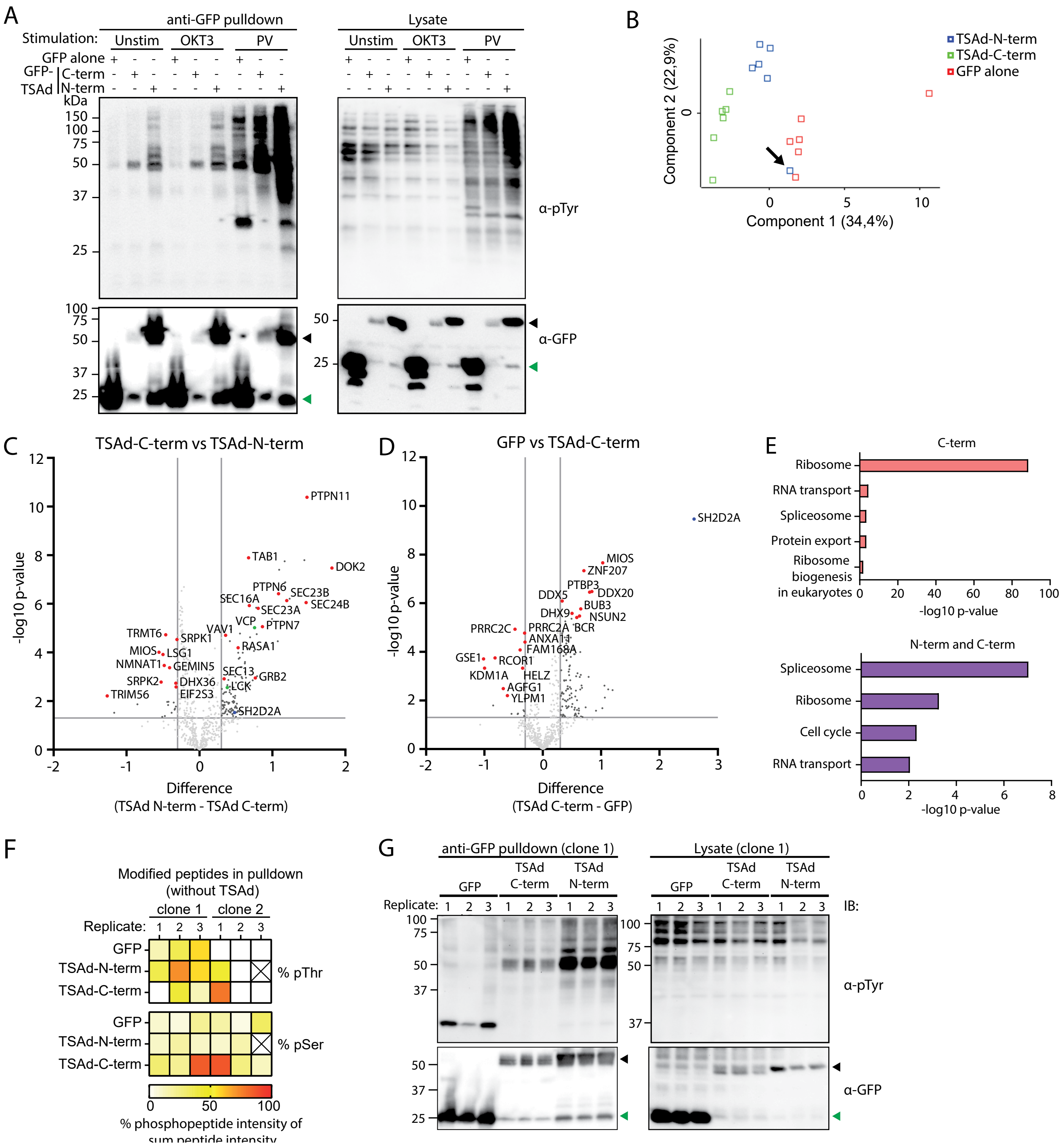


**Supplementary figure 1. AP-MS analysis reveals novel interactors of the TSAd-SH2 domain.** A. JTAg Lck Y192E/Y505F cells were transfected with GFP-tagged TSAd encoding plasmids. 24 hrs later, cells were left unstimulated, treated with anti-CD3e (OKT3) antibody for 2.5 min or treated with pervanadate (PV) for 5 min. Lysates were subjected to GFP pulldown and analysed by immunoblotting. Arrows in black point to TSAd N-term or TSAd C-term GFP tagged constructs, arrow in green point to GFP alone. B. PCA plot of samples ($n$ = 6) following analysis of MS results (after removal of values with less than 5 valid hits). Each point represents a replicate of GFP alone (red), TSAd-C-term (green) or TSAd-N-term (blue). The outlier, which was removed from analysis, is indicated by a black arrow. C-D. Volcano plot of the identified hits in GFP vs TSAd-C-term (D) or TSAd-C-term vs TSAd-N-term by MS analysis. Selected novel interactors marked in red and previously identified ligands marked in green. TSAd (SH2D2A) marked in blue. Horizontal cut-off line denotes p-value = 0.05. Vertical cut-off lines denote fold change = 2. C. Proteins enriched in TSAd-C-term (left) and TSAd-N-term (right). D. Proteins enriched in GFP (left) and TSAd-C-term (right). E. Bar graphs showing enriched KEGG pathways, performed using STRING analysis, with hits identified in Fig 1E. F. Heatmap of phosphopeptides containing phospho-serine (pSer) or phospho-threonine (pThr) in pulldown. % of phosphopeptides intensity is calculated as the ratio between total phosphopeptide intensity vs the total peptide intensity. The "X" in the figure indicates the sample, which was removed as indicated in panel B. G. Immunoblot of GFP-pulldown biological replicate clone 1, subjected to MS analysis in Fig. 1. Arrows in black point to TSAd N-term or TSAd C-term GFP tagged constructs, arrow in green points to GFP alone.

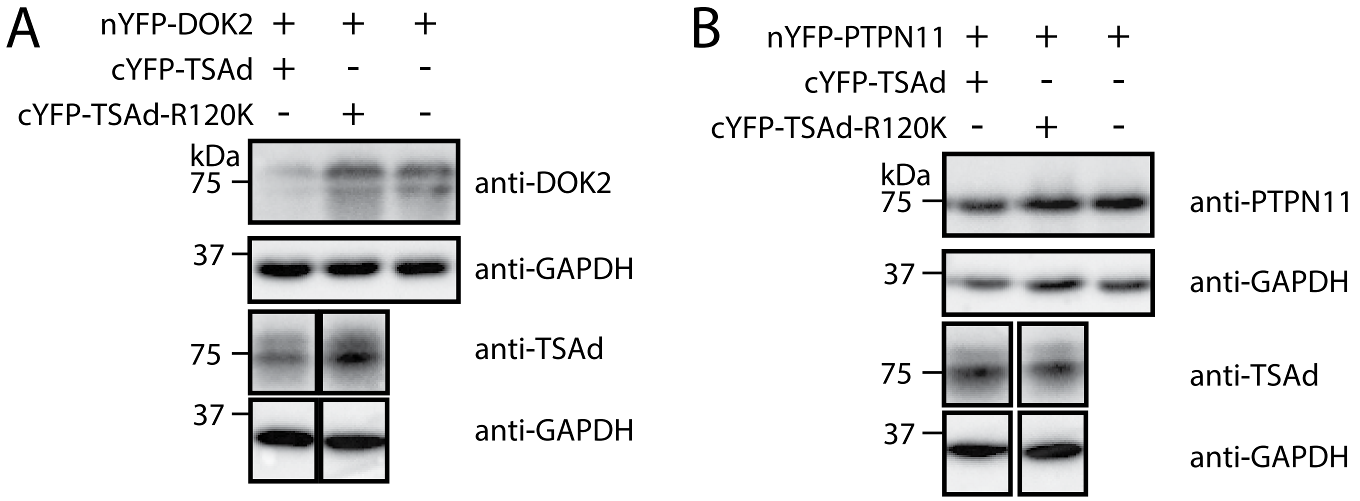


**Supplementary figure 2. Expression of constructs for BiFC assay.** JTAg TSAd KO cells were transfected with plasmids encoding nYFP and cYFP tagged constructs. After 48 hrs, cells were activated with pervanadate for 5 min, washed, and incubated for 4 hrs. Fraction of cells were used for BiFC (Fig. 3) Lysates from cells transfected with DOK2 (A) and PTPN11 (B) were subjected to immunoblotting.

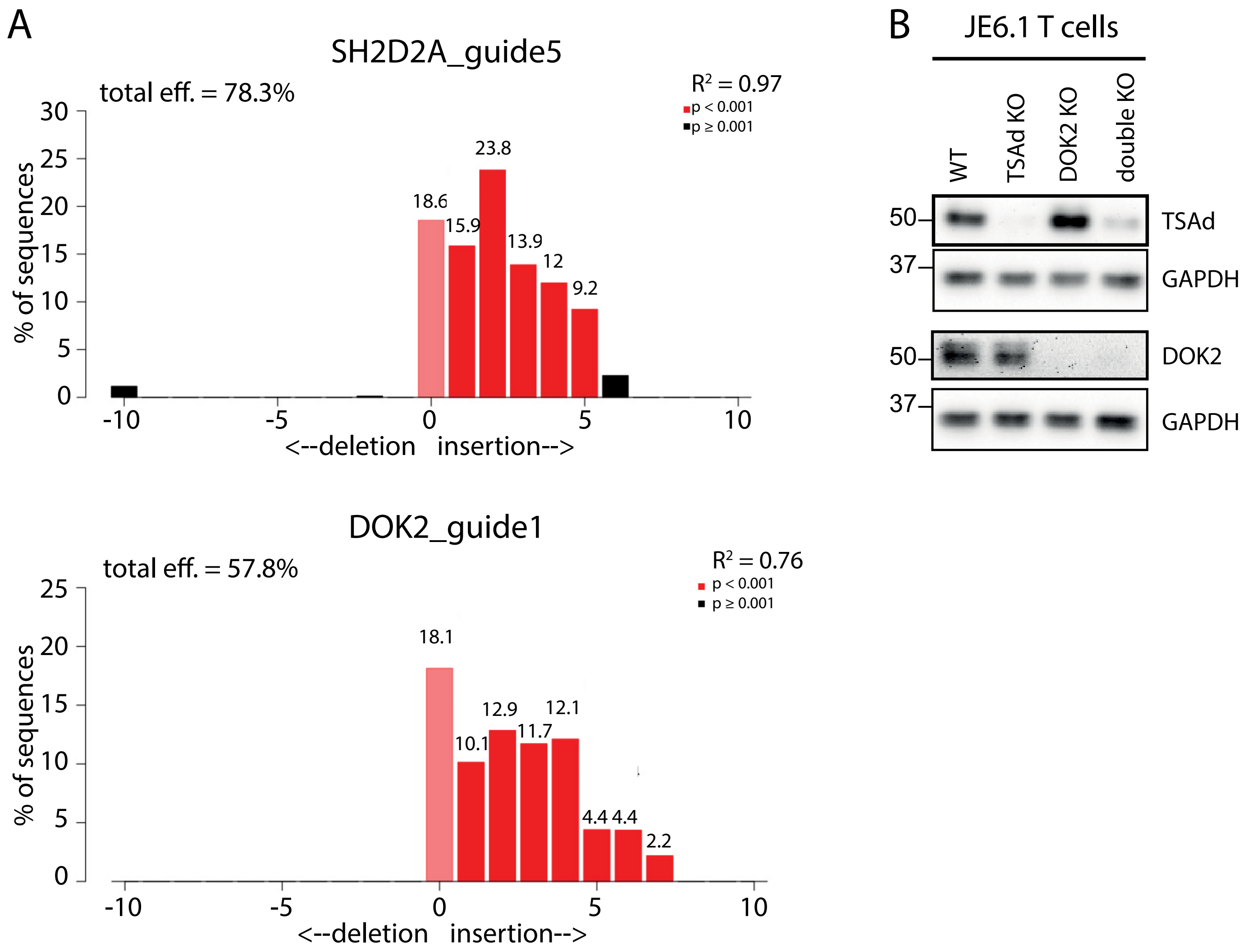


**Supplementary figure 3. CRISPR/Cas9 knockout of TSAd and/or DOK2.** TSAd alone, DOK2 alone or both (double KO) were knocked-out in Jurkat E6.1 cells by CRISPR/Cas9 genome editing. A. Gene editing efficiency was evaluated by TIDE using Sanger sequencing of region of interest. One representative TIDE analysis shown for guides targeting TSAd (top) and DOK2 (bottom). B. One representative immunoblot showing TSAd and DOK2 protein expression in gene edited cells ($n = 2$).
